# Regulatory features aid interpretation of 3’UTR Variants

**DOI:** 10.1101/2023.08.01.551549

**Authors:** Lindsay Romo, Scott D. Findlay, Christopher B. Burge

**Affiliations:** Harvard Medical Genetics Training Program Boston Children’s Hospital Boston, MA 02115; Department of Biology Massachusetts Institute of Technology Cambridge, MA, 02142

**Keywords:** Genetic variants, 3ʹ UTR, RNA-binding proteins, polyadenylation, miRNA, variant interpretation, regulatory motifs

## Abstract

Our ability to determine the clinical impact of variants in 3’ untranslated regions (UTRs) of genes remains poor. We provide a thorough analysis of 3’UTR variants from several datasets. Variants in putative regulatory elements including RNA-binding protein motifs, eCLIP peaks, and microRNA sites are up to 16 times more likely than other variants to have gene expression and phenotype associations. Heterozygous variants in regulatory motifs result in allele-specific protein binding in cell lines and allele-specific gene expression differences in population studies. In addition, variants in shared regions of alternatively polyadenylated isoforms and those proximal to polyA sites are more likely to affect gene expression and phenotype. Finally, pathogenic 3’UTR variants in ClinVar are 20 times more likely than benign variants to fall in a regulatory site. We incorporated these findings into RegVar, a software tool that interprets regulatory elements and annotations for any 3’UTR variant, and predicts whether the variant is likely to affect gene expression or phenotype. This tool will help prioritize variants for experimental studies and identify pathogenic variants in patients.

## INTRODUCTION

Despite the ubiquity of exome sequencing in modern human genetics, our ability to determine the clinical impact of genetic variants remains limited. The average patient has 20,000 variants in their exome, most of which are rare in the population^1^. The effect of rare variants in coding regions can be predicted by their impact on protein amino acid composition^2–4^. However, many variants in the ClinVar variant database are noncoding variants of uncertain significance^5^. The impact of these variants is difficult to predict due to our incomplete knowledge of the function of noncoding regions.

The 3’ untranslated region (3’UTR) comprises the bulk of noncoding sequences present in exomes, and is important for regulation of messenger RNA (mRNA) processing, stability, translation and localization. Sequence-specific RNA-binding proteins (RBPs) interact with cognate RNA motifs at specific 3’UTR positions^6^. Such RBPs often recruit effector proteins to the mRNA that can alter transcript stability, translational efficiency, and intracellular mRNA localization^7^. Altered transcript stability and translation impact protein abundance, while altered transcript localization can impact protein function^7^. Recognition of polyadenylation signals (PAS) within 3’UTRs directs cleavage and polyadenylation of the mRNA transcript. Many 3’UTRs contain more than one functional PAS, and alternative polyadenylation (APA) yields transcripts of different lengths containing different sets of RBP and microRNA (miRNA) target sites^8^. miRNAs are small noncoding RNAs that bind to 3’UTRs when complexed with proteins in miRNPs^9^. Binding is mediated primarily by complementarity between the 3’UTR target and nucleotides 2-7 or 2-8 of the miRNA, called the seed sequence. miRNA binding decreases transcript stability or represses translation^10^.

RBP motifs in the 3’UTR can be predicted from in vitro binding studies using RNA Bind-n-Seq (RBNS), RNACompete or other assays, or identified in cells via enhanced crosslinking and immunoprecipitation (eCLIP)^11, 12^. miRNA sites can be predicted in silico from base complementarity and sequence conservation, or identified in cells using crosslinking and sequencing^13, 14^. Variants in the 3’UTR, especially those that disrupt RBP interactions, miRNA binding, or cleavage and polyadenylation, are likely to impact mRNA function and may be deleterious, potentially contributing to disease.

Several metrics have been developed to predict the pathogenicity of noncoding variants^15–20^. However, these metrics don’t take into account the unique regulatory features of 3’UTRs, such as alternative polyadenylation, RBP interactions, or miRNA binding. More general methods that may implicate 3’UTR variants include expression quantitative trait loci (eQTLs), which link variants to gene expression changes, and genome wide association studies (GWAS), which link variants to phenotypes^21–24^. However, these methods don’t suggest mechanisms, detect association rather than causality, and can only interrogate common variants. Experimental methods such as massively parallel 3’UTR reporter assays can address molecular functions of variants, with the caveats that variants are assayed in artificial genomic contexts and are over-expressed^25, 26^. Saturation genome editing can generate possible genomic variants in clinically-important regions, with clonal cell lines used to assess phenotypes^27^. Though effective in identifying pathogenic variants, this method is laborious and expensive.

Exome and genome sequencing are becoming less expensive and more rapid, but data interpretation remains the limiting step. For many patients, exome or genome sequencing results in a diagnosis that alters clinical management and can be life-saving^28, 29^. More general and accessible methods of variant characterization could therefore be highly impactful. We propose that 3’UTR variants that disrupt (or create) specific types of regulatory elements are more likely to alter function and contribute to disease, aiding in the identification and interpretation of pathogenic 3’UTR variants.

Here, we show that single nucleotide variants (SNVs) in locations that overlap RBP motifs, eCLIP peaks, and miRNA sites are both more evolutionarily conserved and more likely than other variants to be associated with phenotypic or gene expression changes. Many of these variants are causal, as they are enriched for allele-specific RBP binding in cell lines, and allele-specific gene expression differences in population studies. eQTL variants in potential regulatory elements are more likely to be GWAS hits, and pathogenic 3’UTR variants in ClinVar are more likely to fall in a regulatory element than benign variants. We also show that variants in certain 3’UTR regions, e.g., proximal to polyA sites, are more likely to be linked to gene expression changes or phenotypes. Finally, we provide a high-throughput R package, RegVar, which assesses regulatory elements and annotations associated with any 3’UTR variant of interest, and predicts whether the variant is likely to affect gene expression or phenotype. To our knowledge, this is the first program to specifically characterize 3’UTR variants. We expect the program will help prioritize variants for experimental studies and identify thousands of pathogenic variants.

## MATERIALS AND METHODS

### eQTL processing

DAGP fine-mapped eQTL variant call files were downloaded from the GTex project (https://gtexportal.org/home/datasets) and intersected with terminal 3’UTRs using BEDTools, as defined by the region from the GENCODE stop codon to the most distal polyA Database peak^24, 30–32^. Because many variants in 3’UTRs (and elsewhere) are in linkage disequilibrium, it can be difficult to discern causal variants. Fine-mapping is a statistical method to distinguish the effects of variants in linkage disequilibrium blocks^33^. Fine-mapping eQTLs or GWAS hits results in a posterior inclusion probability (PIP) for each variant that represents the likelihood each is causal of expression differences or phenotypes^34^. For each variant-tissue combination, only the tissue with the highest PIP was used, except for the transcript expression analysis (see below).

### GWAS processing

SUSIE fine-mapped UK BioBank GWAS variant call files were downloaded from the Finucane lab website (https://www.finucanelab.org/data) and intersected with 3’UTRs, as above^35^. All variant-phenotype combinations were considered in analysis.

### Identifying variants in putative 3’UTR regulatory elements

3’UTRs were defined as above. To identify variants in putative RBP motifs, reference/alternative variants and their surrounding genomic sequence were processed with RBPamp (ref). Variants overlapping an RBNS motif with an affinity of >0.33 of the ideal motif were considered to be in RBP motifs. Alternate variants overlapping a motif with an affinity of 0.66 or more compared to the ideal motif were considered preserving, and alternate variants that caused the motif affinity to drop below 0.33 compared to the ideal were considered disrupting. To identify variants in eCLIP peaks, variants were intersected with eCLIP coordinates downloaded from ENCODE (https://www.encodeproject.org/)^12^. For Figure 3A, we considered RBPs with at least fifty variants in peaks to allow sufficient power to detect PIP differences. RBPamp eCLIP- Proximal (ReP) sites were defined as motifs matching the highest affinity RBPamp motif in the vicinity of each of the eCLIP peaks. Variants in possible conserved family miRNA sites and their seed types and site conservation were defined by TargetScan^13^.

### Annotation of 3’UTR variants

PolyA signals/sites and 3’UTR isoforms were identified from aggregate 3Pseq data generated from multiple tissues and cell lines^36^. Calling of polyA sites and removal of peaks from genomic polyA priming was performed on 3PSeq data as described previously^37^. We considered variants proximal to polyA sites if they fell within 50 nucleotides of a 3Pseq peak. The relationship between distance to polyA site and PIP becomes nonsignificant in regression models for variants more than 50 nucleotides away. Shared regions of APA isoforms were regions proximal to the first polyA site. Partially-shared regions were between the first and penultimate polyA site, whereas unique regions were distal to the penultimate polyA site.

### Conservation

Variants were merged with conservation information from PhastCons 100-way scores to identify conserved (PhastCons>0.5) or non-conserved (PhastCons<0.5) variants^38^. For miRNA sites, the TargetScan conservation designation was used^13^.

### Statistical analysis

The proportion of causal variants in an element (the fraction with a PIP of greater than 0.25) was calculated using pairwise proportion z-tests. The proportions causal for variants in elements was compared to that for variants not in elements using a Fischer exact test. When comparing PIPs, a paired Wilcoxon rank sum test was used. For odds ratios, a Fischer exact test was used to determine confidence intervals and significance. For the generalized linear model, we used a binomial distribution to model a binary score (GWAS or eQTL PIP greater or less than/equal to 0.5). Goodness of fit was assessed via Hosmer-Lemeshow Test. All p-values were corrected using the false discovery rate method.

### Expression analysis

The relative expression of transcripts with the alternate versus reference allele was determined by the transcript normalized effect size (NES) from GTex^24^. The NES is the slope of the eQTL regression line comparing expression of transcripts with the alternative and reference alleles; more positive NES values indicate higher gene expression in individuals with the alternative allele, and vice versa^24^. Each PIP-tissue combination was considered for every eQTL.

### BEAPR analysis

All heterozygous variants in eCLIP peaks in K562 and HepG2 cells, as well as their predicted eCLIP allele-specificity as defined by BEAPR analysis, were kindly provided by the Xiao lab^39^. Only 3’UTR variants were considered, and variants in RBP motifs (with reference allele affinity >0.05 of alternate allele) matching the eCLIP RBPs (defined as above) were compared to variants only in eCLIP peaks but not in motifs.

### ClinVar analysis

Variant call files were downloaded from ClinVar (https://www.ncbi.nlm.nih.gov/clinvar/) and intersected with 3’UTR coordinates/annotations and putative regulatory elements as above^5^. Only variants with known clinical impact (pathogenic or benign) were considered.

### CADD scores

Raw as well as scaled CADD scores for all gnomAD variants were downloaded from the CADD website (https://cadd.gs.washington.edu/) and intersected with eQTL variants^15, 40^. The raw CADD scores of variants in putative regulatory elements were compared to those for controls.

### RegVar

The development version of the RegVar R package is available for download on github at https://github.com/RomoL2/RegVar. The RegVar tool characterizes user-provided 3’UTR variants by their regulatory features as described. A variant is predicted to be an eQTL or GWAS hit if its log-odds is greater than 0.01 (eQTL) or 0.0075 (GWAS) using our logistic regression model (see ‘statistical analysis’). These thresholds maximize sensitivity and specificity.

## RESULTS

### Identification of causal 3’UTR variants

We developed an analysis pipeline to identify features that might differentiate 3’UTR variants that impact gene expression or phenotype (Figure 1). We used three sources of 3’UTR variants: fine-mapped eQTLs identified by GTEx (82,903 variants), fine-mapped GWAS hits from the UK Biobank (174,065 variants), and heterozygous variants in eCLIP peaks (2,856 variants)^22, 24, 39^. Fine-mapping is a statistical method that yields a posterior inclusion probability (PIP) for each eQTL or GWAS hit representing the likelihood that each is causal for the observed association^33, 34^.

**Figure 1:**
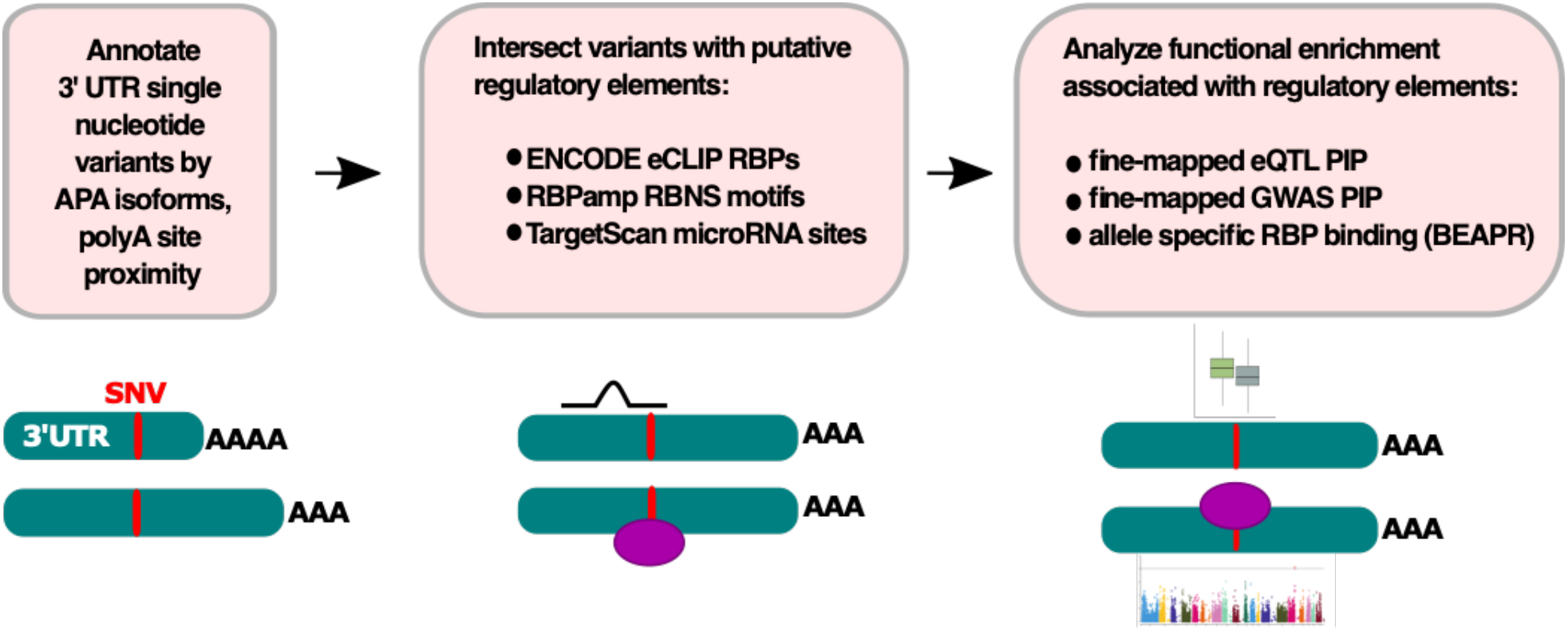
Variant processing pipeline schematic. We compared the probability that eQTLs and GWAS hits affect gene expression (eQTLs) or phenotype (GWAS hits) for variants in eCLIP peaks, RBP motifs, and various 3’UTR annotation categories. We compared the likelihood of overlapping an RBP motif for heterozygous variants with and without allele-specific eCLIP binding.

Variants were first annotated by APA isoform location, and then intersected with putative regulatory elements. The specific categories of elements studied included RBP binding motifs from in vitro studies, eCLIP peaks, “ReP” sites, and miRNA sites (see methods)^11–13, 36^. Here, we defined “ReP” (RBPamp eCLIP-Proximal) sites for RBPs with both in vitro and in vivo binding data as the highest-affinity motif for the RBP in the vicinity of each of its eCLIP peaks^11, 12^. To determine whether variants in specific regulatory element or annotation categories preferentially impact gene expression or phenotype, we compared the PIP for variants located in these elements versus controls. Allele-specific eCLIP binding events^39^ were used to assess whether variants in motifs altered RBP binding.

### 3’UTR variants in putative regulatory elements are associated with altered gene expression

We hypothesized that variants in RBP motifs and/or eCLIP peaks often impact transcript expression by altering binding of regulatory RBPs. Overall, most eQTL variants have low fine-mapped PIP values (Figure S1). However, we found that high-PIP eQTLs are slightly more likely than non-eQTLs to be located in an in vitro-derived RBP motif, over four times more likely to be in an eCLIP peak, and over nine times more likely to be in a ReP site (Figure 2A). These observations support our hypothesis that each of these classes of elements is enriched for variants that alter expression. Variants located in eCLIP peaks and ReP sites also have significantly higher PhastCons 100-way conservation scores than controls, even after matching for gene expression level, providing further evidence that these classes are enriched for regulatory function (Figure 2B)^41^.

**Figure 2:**
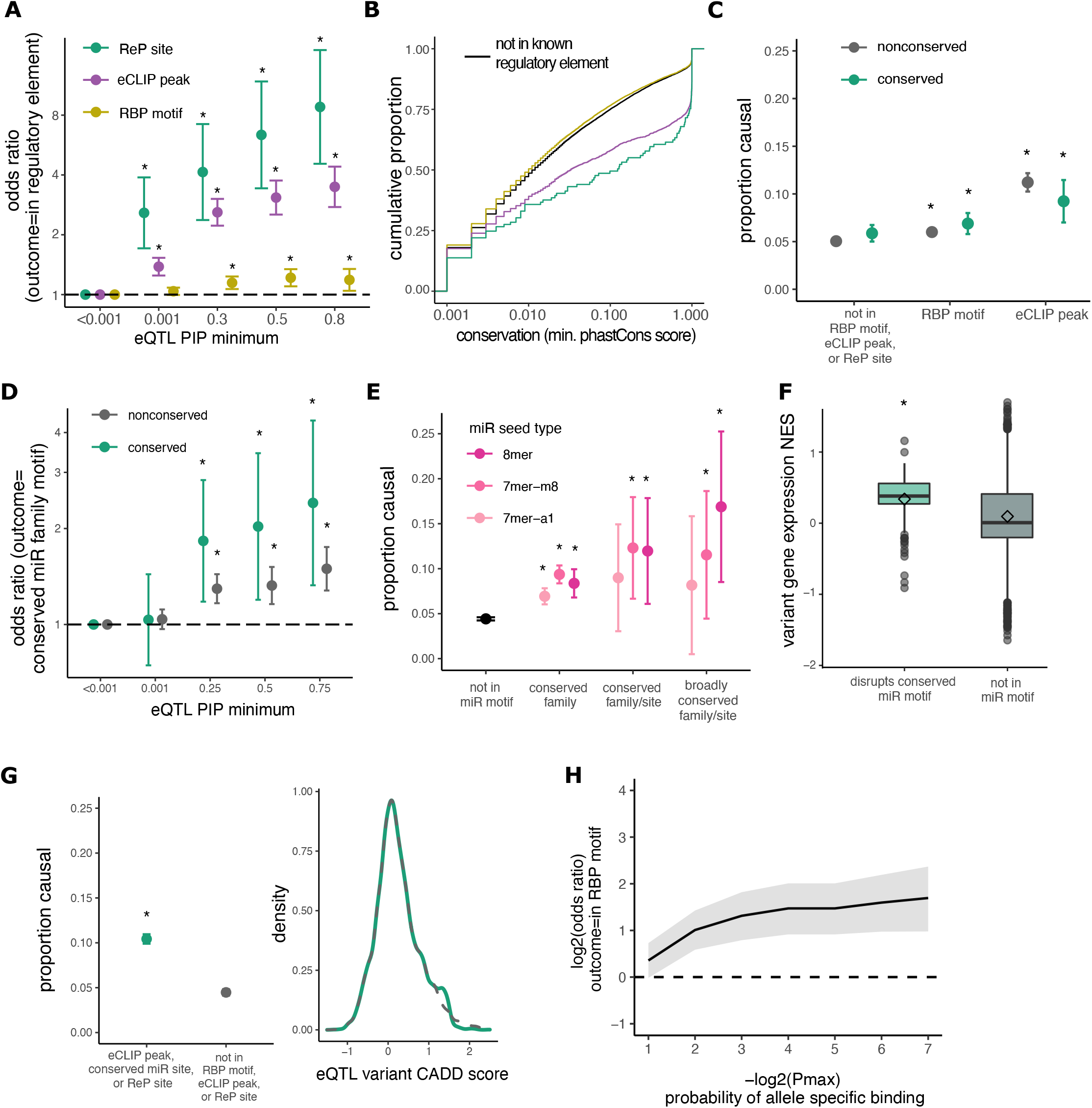
3’UTR variants in RBP motifs, eCLIP peaks, and miRNA sites are associated with gene expression changes. **A** Odds of an eQTL variant being in predicted RBP sites versus control variants (PIP<0.001) as minimum PIP increases; odds ratio is shown with 95% confidence intervals; *p<0.01. **B** Comparison of PhastCons score distributions for eQTL variants in ReP sites (green), eCLIP peaks (purple), RBP motifs (orange), or outside of known regulatory elements (black). **C** Proportion causal (PIP>0.25) for eQTL variants not in RBP motifs or eCLIP peaks compared to variants in RBP motifs or eCLIP peaks. **D** Odds of an eQTL variant being in predicted miRNA site versus control variants (PIP<0.001) as minimum PIP increases, as in A. **E** Proportion causal for variants not in miRNA sites compared to variants in miRNA sites of different seed types. **F** Variant NES (from GTEx) on gene expression for high-confidence (PIP>0.9) eQTLs disrupting miR motifs (green), or not in RBP/miR motifs or eCLIP peaks (gray). **G** Proportion causal (left) for variants not in RBP motifs or eCLIP peaks compared to variants in ReP sites or eCLIP peaks, matched for raw CADD score (right). **H** Odds of heterozygous variants in HepG2 and K562 cells being in an RBP motif for decreasing allele-specific eCLIP binding p-values, with 95% confidence intervals.

We considered the proportion of eQTL variants with PIP > 0.25 (i.e. variants at least 25% likely to alter expression) as a summary statistic, which we call “proportion causal”. Although the precise PIP cutoff used is somewhat arbitrary, repeating our key analyses using different PIP thresholds yielded similar results (Figure S2). We found that eQTLs in RBP motifs and those in eCLIP peaks had a higher proportion causal than variants outside of these elements, and that variants in RBP motifs were more likely to be causal if conserved (Figure 2C). Unfortunately, the number of eQTL variants in ReP sites was too low to perform a similar analysis.

We hypothesized that variants in miRNA target sites typically disrupt miRNA binding, resulting in increased mRNA levels. Indeed, high-PIP eQTLs are over twice as likely to fall in conserved miRNA sites than non-eQTL variants (Figure 2D). miRNAs with greater complementarity to target transcripts (longer seed matches: 8mer > 7mer-m8 > 7mer-a1) exert stronger regulatory effects^13^. We found that eQTLs in conserved 8mer sites of miRNAs in broadly conserved families have three times higher proportion causal than controls (Figure 2E).

### 3’UTR variants in putative regulatory elements likely cause expression changes

To determine whether variants in miRNA sites increase gene expression, as expected, we compared the normalized effect size (NES) of causal (PIP>0.25) eQTLs that disrupt miRNA motifs to those outside of predicted regulatory elements. The NES measures the magnitude and direction in which eQTLs change gene expression^24^. Variants predicted to disrupt conserved miRNA motifs predominantly have positive NES (P < 0.001, Wilcoxon rank-sum test), suggesting a direct relationship between disrupted miRNA binding and increased expression (Figure 2F).

If variants in RBP motifs and eCLIP peaks commonly alter gene expression, we would expect some to be pathogenic. The Combined Annotation Dependent Depletion (CADD) score is a metric that discriminates between benign and pathogenic variants based on their evolutionary deleteriousness. We found that variants in eCLIP peaks and RBP motifs have significantly higher CADD scores (Figure S3). As CADD score does not incorporate RBP motif or eCLIP information, higher scores reflect other features of RBP motifs such as conservation and base composition. We found that variants in eCLIP peaks, conserved miRNA sites, or ReP sites are more likely to cause gene expression changes than controls, even when comparing sets with matched CADD scores (Figure 2G). Thus, considering eCLIP, miRNA, and ReP information can add substantially to the information in CADD scores for identification of functional variants.

To test the idea that expression differences associated with eQTLs in RBP motifs result from differential binding of RBPs, we analyzed allele-specific eCLIP binding^39^. Heterozygous variants that exhibit allele-specific eCLIP enrichment are up to four times more likely to be located in a motif for the corresponding RBP than variants that do not (Figure 2H). This observation supports that the gene expression changes associated with variants in eCLIP peaks noted above commonly result from changes in RBP binding.

### Enrichment for RBP motifs and eCLIP peaks amongst eQTLs is driven by a subset of RBPs

We next examined which RBPs are responsible for increased PIPs among eQTLs in eCLIP peaks. RBPs with high-PIP eQTLs in eCLIP peaks included those known to bind the 3’UTR to alter mRNA stability, such as PABC4 and LARP4^42, 43^. In contrast, RBPs with fewer high-PIP eQTLs in eCLIP peaks included transcription factors or repressors such as BCLAF1 as well as primarily nuclear proteins such as HNRPL, KHSRP, and QKI (Figure 3A). Mean PIP values were positively correlated across RBPs for variants in RBP motifs and those in eCLIP peaks (Figure 3B), suggesting that these two subsets of variants function similarly, with some RBPs impacting expression more often than others.

**Figure 3:**
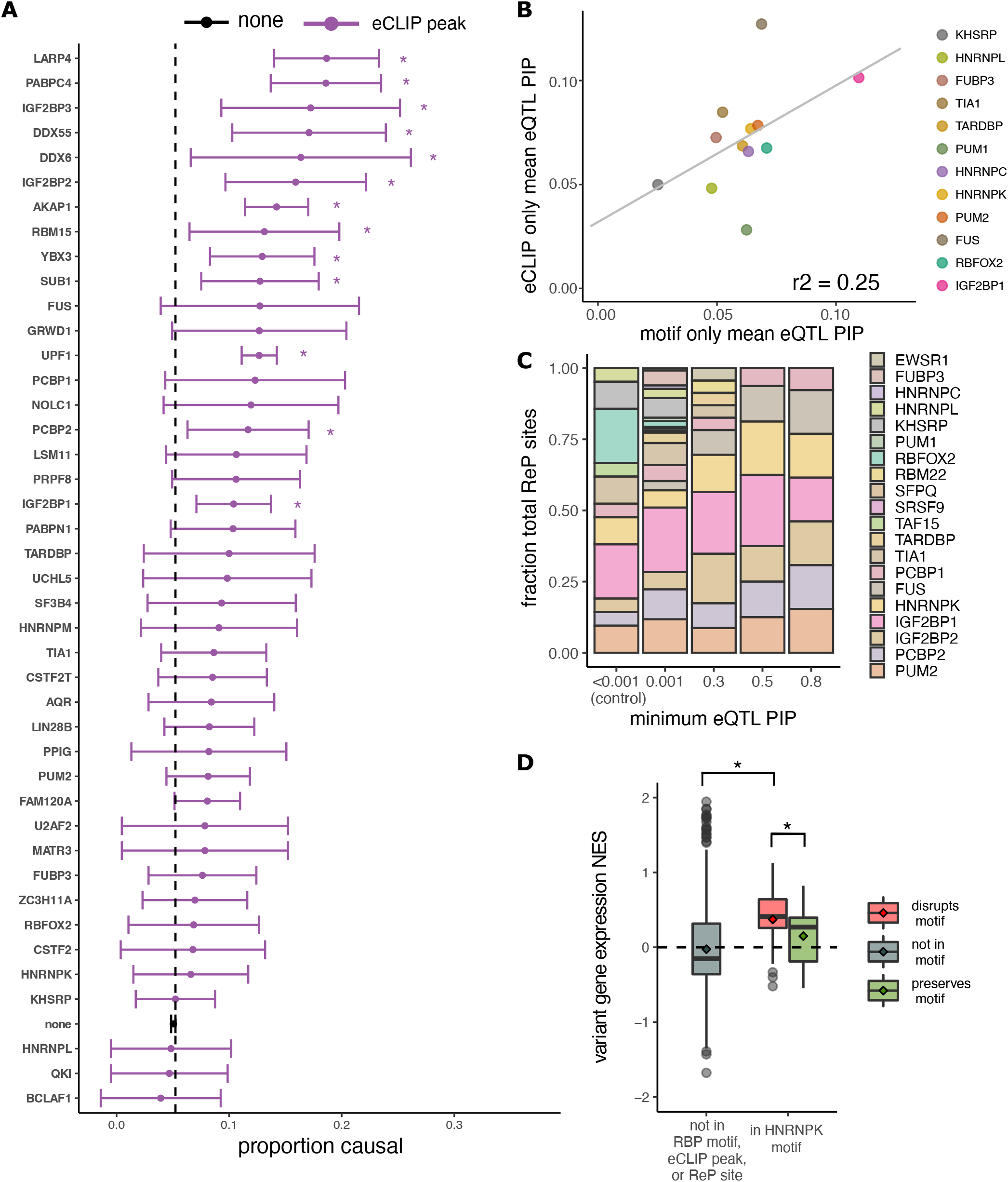
Certain RBP motifs and eCLIP peaks are enriched amongst eQTLs and alter expression. **A** Proportion causal (with 95% confidence interval) for variants in eCLIP peaks for different RBPs, for all RBPs with ≥ 50 variants in eCLIP peaks). The proportion causal for variants not in any eCLIP peak was 0.05 (dashed line). * Indicates P<0.05. **B** Mean PIP for eQTLs in eCLIP peaks but not RBP motifs (y-axis) versus mean PIP for eQTLs in RBP motifs but not eCLIP peaks, for all RBPs in both datasets. Shown is the regression line with Pearson correlation coefficient. **C** Distribution of RBPs amongst ReP sites at different minimum eQTL PIP cutoffs. Shown are RBPs representing at least 0.1% of all ReP sites. **D** Variant normalized effect size (NES, from GTEx) on gene expression for high-confidence (PIP>0.9) eQTLs not in RBP motifs or eCLIP peaks (gray), and for high-confidence eQTLs predicted to disrupt (red) or preserve (green) HNRNPK motifs.

To further explore which RBPs may most commonly impact expression, we examined ReP sites amongst high-PIP eQTLs and found that several motifs, including those for HNRNPK, are highly represented (Figure 3C). HNRNPK is a multifunctional RBP involved in both transcriptional and post-transcriptional mRNA processing that binds 3’UTRs at C-rich motifs to alter stability of target mRNAs^44–46^. We found that high-confidence eQTLs that disrupt HNRNPK motifs are associated with higher transcript expression than those that preserve motifs or are located outside of HNRNPK motifs (Figure 3D). This observation suggests that most HNRNPK binding tends to destabilize mRNAs in tissues assessed for eQTLs.

### 3’UTR variants in putative regulatory elements likely result in phenotypic changes

Recent studies have demonstrated limited overlap between GWAS hits and eQTLs, and found that many genes with eQTLs are under weak selective constraint and are likely less functionally important than genes with GWAS hits^47^. Therefore, it was of interest to explore the extent to which eQTL variants that change gene expression by disruption of RBP motifs have phenotypes. To assess the effect of 3’UTR variants on phenotypes, we analyzed GWAS data generated by the UK BioBank^22^. Overall, GWAS variants have lower PIPs than eQTL variants after fine-mapping. Otherwise, the distribution of GWAS and eQTL variants along the 3’UTR was similar and uniform (Figure S1).

GWAS hits in eCLIP peaks had much higher conservation scores than those outside of eCLIP peaks, even those in an RBP motif (Figure 4A). To determine whether variants in regulatory elements result in phenotype changes, we compared PIPs for variants in RBP motifs, eCLIP peaks, and controls. GWAS hits in these elements have higher PIPs compared to variants outside of these sites (Figure 4B). (Too few ReP sites overlapped to permit similar analyses of this class.) Conservation has a larger impact on GWAS variant PIP than on eQTL PIP (Figure 4B), likely because variants that affect phenotype are under stronger selection than those that merely affect gene expression, consistent with recent studies comparing GWAS hits and eQTL variants^47^. Also potentially contributing to observed differences in PIP is the fact that most GWAS traits are categorical whereas eQTLs are continuous.

**Figure 4:**
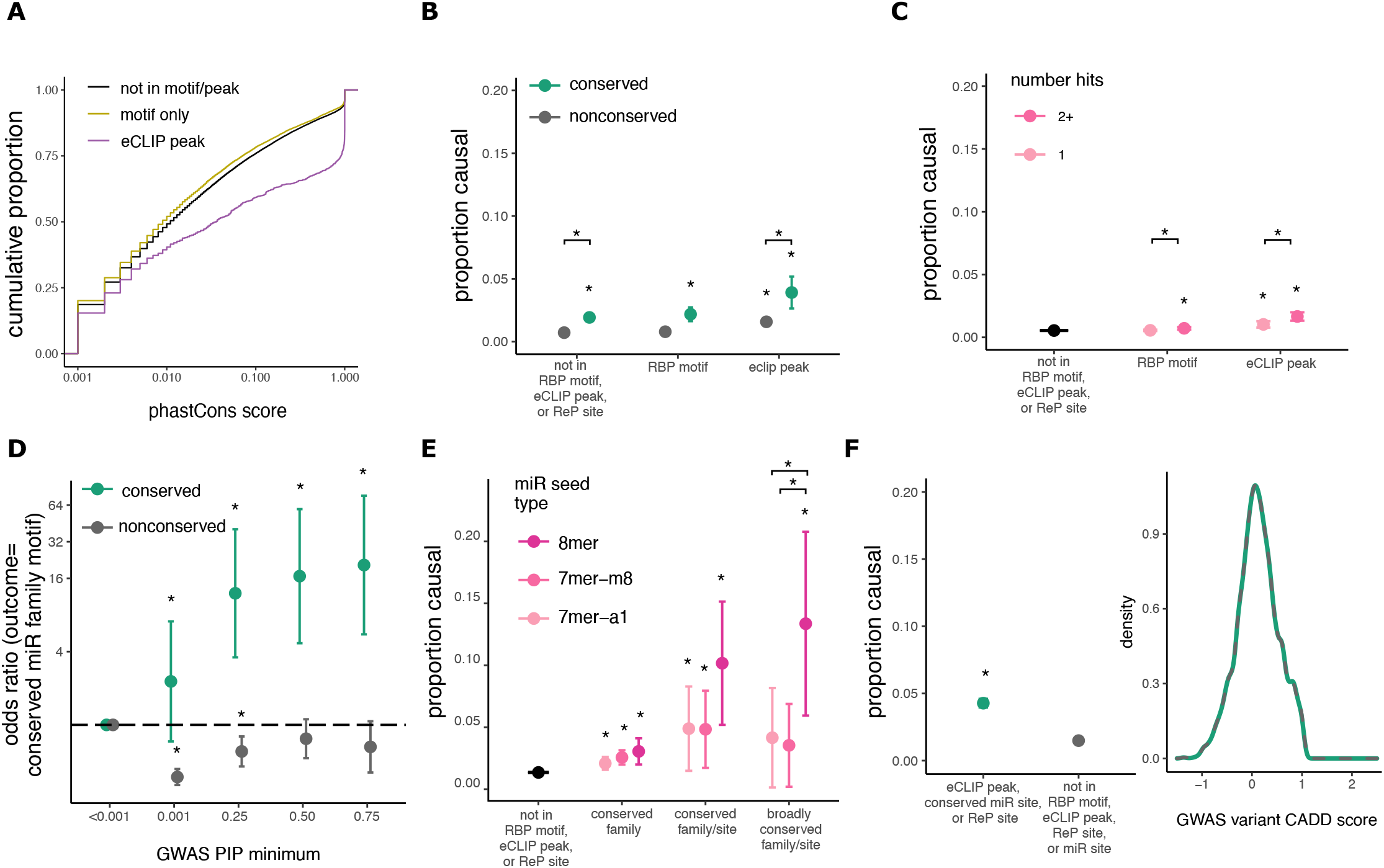
3’UTR variants in RBP motifs, eCLIP peaks, and miRNA sites are associated with phenotypes. **A.** Comparison of PhastCons score distributions for GWAS variants in eCLIP peaks, RBP motifs, or no known regulatory elements. **B** Fraction causal (proportion of GWAS hits with PIP>0.25) for variants not in RBP motifs or eCLIP peaks compared to variants in RBP motifs or eCLIP peaks. * Indicates P<0.05. **C** As in B, but for variants in a single motif or CLIP peak compared to variants in more than one motif or peak in genes matched by gene expression. **D** Odds of a GWAS variant being in predicted miRNA site versus control variants (PIP<0.001) as minimum PIP increases; shown is odds ratio with 95% confidence intervals. **E** Fraction causal for variants not in miRNA sites compared to variants in miRNA sites with increasing predicted seed strength. **F** Proportion causal (left) for variants not in RBP motifs or eCLIP peaks compared to variants in ReP sites or eCLIP peaks, matched for raw CADD score (right).

Recent research has suggested that variants in “RBP hubs” – locations where multiple RBPs bind – have lower allele frequencies compared to variants in single eCLIP peaks. However, these studies did not control for the bias in eCLIP data toward genes with higher expression, a property which is associated with higher conservation^48^. Regardless, we found that variants located in multiple overlapping eCLIP peaks or RBP motifs are slightly more likely to be causal of phenotypes than those in single peaks/motifs, even after controlling for gene expression (Figure 4C), supporting that such variants are enriched for function. Binding of multiple RBPs to a region may increase the likelihood that the variant alters binding of at least one protein, or may enrich for sites that have multiple or important functions.

Variants in miRNA sites are also enriched for association with GWAS phenotypes. As PIP increases, GWAS variants are up to sixteen times more likely to be located in conserved miRNA target sites. However, GWAS variants are actually mildly depleted from non-conserved miRNA sites, suggesting that such sites rarely impact phenotype (Figure 4D). Similar to eQTLs, GWAS hits in miRNA sites have higher PIPs, especially those in 8mer seeds, which have up to 15-fold higher proportion of causal variants than variants outside of regulatory elements (Figure 4E). We did not see a significant association between the number of distinct miRNA family targets overlapping a variant and the proportion of high-PIP GWAS variants, but statistical power was limited (Figure S4). Concerns have been raised regarding the accuracy of computational methods for predicting miRNA sites, and the ability of these predicted sites to impact phenotype^49^. Our results suggest that computationally predicted miRNA sites, especially conserved targets for conserved miRNA families, are strongly enriched for causal variants affecting both gene expression and phenotype.

As observed for eQTLs, we found that GWAS hits in eCLIP peaks and RBP motifs have significantly higher CADD scores (Figure S3). Variants in eCLIP peaks, conserved miRNA sites, or ReP sites are more likely to be causal, even when CADD scores are matched, again supporting the argument for supplementing CADD scoring with regulatory information to improve discrimination of pathogenic variants (Figure 4F).

### Effects of 3’UTR variants on expression and phenotype depend on APA isoforms

APA impacts post-transcriptional regulation and steady-state transcript expression^8^. We found that the proportion causal was two-fold higher for eQTLs proximal to polyA sites (within 50 nt) than for eQTLs further from the PAS, for major PAS categories (Figure 5A). This enrichment in a region where core PAS motifs are located likely impacts expression via changing the location or efficiency of cleavage and polyadenylation^8^.

**Figure 5:**
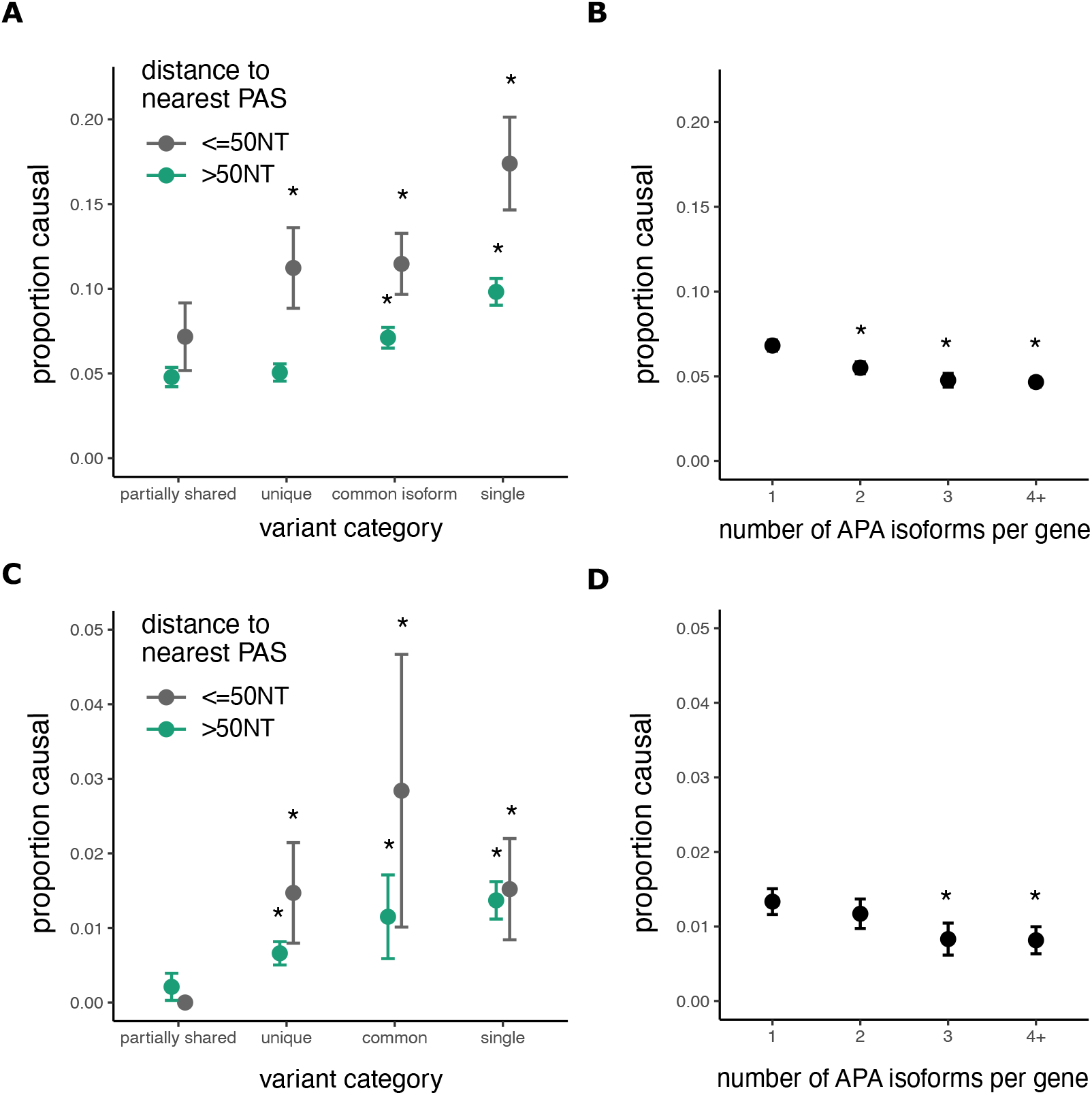
Variants in single or common 3’UTR regions more often impact gene expression and phenotype. **A** Fraction causal (proportion of eQTL variants with PIP greater than 0.25) for variants in various 3’UTR regions and proximal (<50 nucleotides) or distal to polyA sites. * indicates P<0.05 compared to left-most group. **B** Proportion causal for eQTL variants in genes with various numbers of canonical APA isoforms. **C** Proportion causal for GWAS variants in various 3’UTR regions, as in A. **D** Proportion causal for GWAS variants in genes with various numbers of APA isoforms, as in B.

We also observed that the PIPs of eQTLs are higher in genes with single 3’UTR isoforms and in common regions of 3’UTRs from APA genes than in “partially shared” (alternative) UTR regions of the same genes (Figure 5A). The likely explanation is that presence in all transcripts from a gene versus only some transcripts increases the magnitude of the impact on gene expression for common types of variants. Considering the relationship between eQTL PIP and the number of APA isoforms, we found that variants in genes with fewer APA isoforms have a higher proportion of causal variants, likely for similar reasons (Figure 5B). These findings persisted after controlling for 3’UTR length and distance to the stop codon, and the number of eQTL variants per gene did not vary substantially with APA isoform number (Figure S5). Unlike eQTLs, GWAS hits did not have higher PIPs near polyA sites; however, we do see a trend towards higher PIP for GWAS variants in common regions or in 3’UTRs with a single PAS (Figure 5C). Variants in genes with fewer APA isoforms also have higher GWAS PIPs, as seen for eQTLs (Figure 5D).

### Regulatory features help to identify pathogenic 3’UTR variants

Here, we have shown that conserved variants in RBP and miRNA sites within common 3’UTR regions of genes with fewer APA isoforms are more likely to impact gene expression and phenotype. These features can be incorporated into a generalized linear model to predict whether a variant is an eQTL or GWAS hit based on its regulatory features (Figure S6). The model coefficients for each feature represent the increase in the odds of a variant being a causal GWAS hit or causal eQTL (with PIP>0.5) if the variant overlaps the feature. As expected, conservation score has the largest predictive value for GWAS hits, whereas eCLIP peak overlap is most predictive for eQTLs (Figure 6A). Similar to recent studies, we found limited overlap between eQTLs and GWAS hits^47^. For our variants, only 41% of eQTLs were GWAS hits, suggesting that most eQTLs do not impact assayed phenotypes. However, we found that 47% of eQTLs in eCLIP peaks or conserved miRNA sites were GWAS hits, versus 39% of eQTLs located outside of predicted regulatory elements (Figure 6B, P<0.005). Thus, the 3’UTR regulatory features considered here can help to predict which eQTLs are likely to affect phenotype.

**Figure 6:**
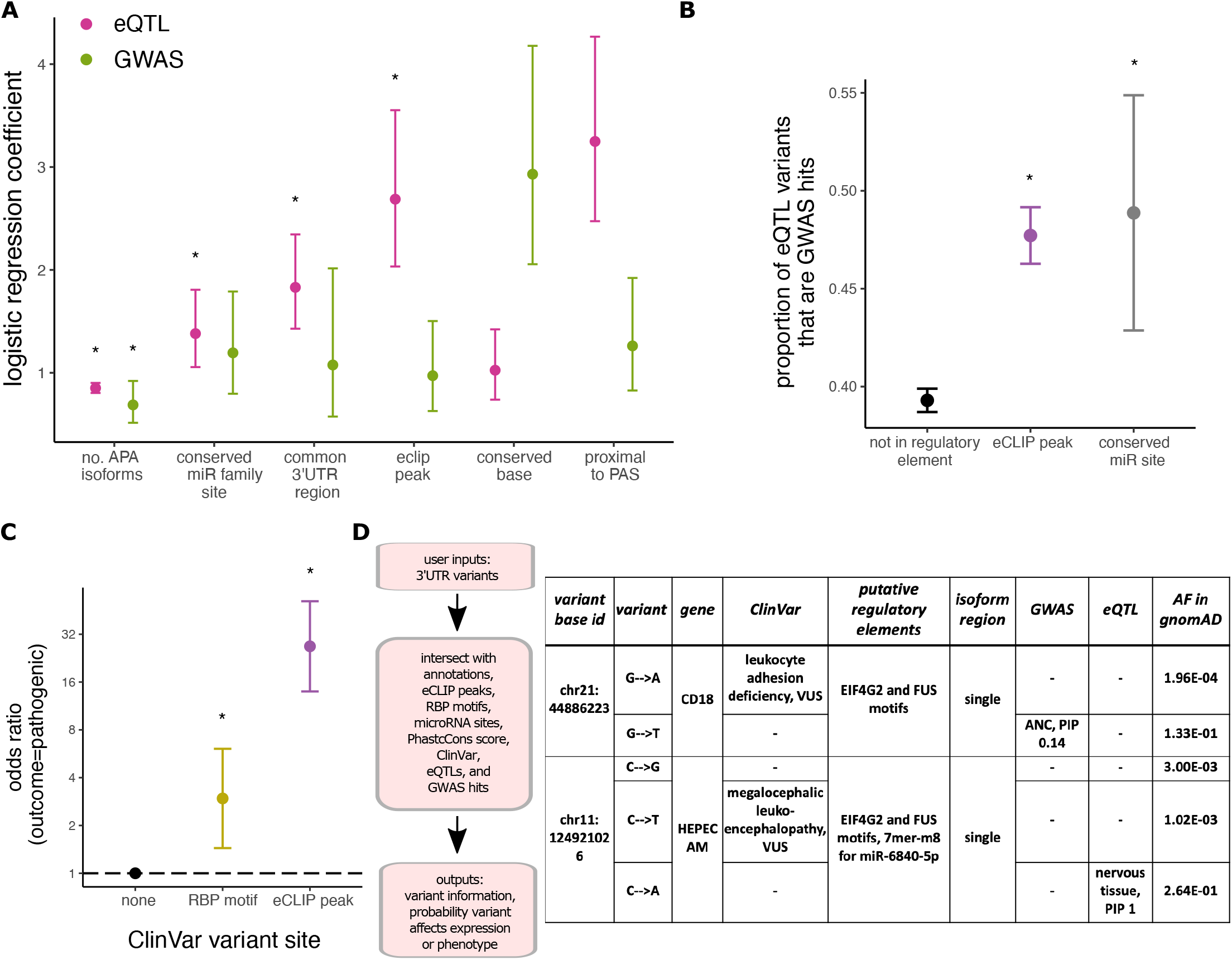
Characterization of 3’UTR variants into their annotations and regulatory elements helps prioritize variants for functional analysis and disease classification. **A** Exponential of logistic regression model coefficients with 95% confidence intervals. The model independent variable is binary (PIP greater or less than/equal to 0.5). * indicates P<0.05. **B**Intersection of eQTL variants in different putative regulatory elements with GWAS hits. **C** Odds of a ClinVar 3’UTR variant in RBP motifs or eCLIP peaks being pathogenic versus variants not in a predicted regulatory element; shown is odds ratio with 95% confidence intervals. **D** RegVar workflow and example output for two ClinVar 3’UTR variants of uncertain clinical significance (genomic coordinates are in hg38).

We show that regulatory analysis of noncoding variants using several orthogonal methods aids in identification of causal eQTLs and GWAS hits, many of which are expected to be pathogenic. Of conserved 3’UTR variants with known clinical significance in the ClinVar database, variants in RBP motifs are 3 times more likely and variants in eCLIP peaks are over 20 times more likely than variants not in regulatory elements to be pathogenic (Figure 6C). (Too few ClinVar variants were located in conserved miRNA sites to permit analysis.) Conservation had little if any impact on these findings as the degree of conservation between these ClinVar variant subsets was similar (Figure S7).

Our results indicate that regulatory analysis of 3’UTR variants can aid in prioritization of variants for functional analysis and detection of pathogenic variants in patients. To this end, we developed a program, RegVar, that characterizes 3’UTR variants by their annotations, conservation, and predicted regulatory elements (Figure 6D, left). This tool will enable prioritization of 3’UTR variants for functional analysis, potentially contributing to improved patient diagnosis and treatment. Many variants in ClinVar are located in untranslated regions, and most of these variants are of uncertain significance^5^. Prioritization of specific variants for experimental analysis will be beneficial to the many patients who have no genetic diagnosis despite whole exome or whole genome sequencing.

Variants causing disease typically have low allele frequency in the general population as a result of natural selection, whereas the GWAS and eQTL analyses have statistical power only to detect variants of high frequency. As expected, we found little to no overlap between eQTL and GWAS 3’UTR variants and ClinVar variants. However, as a sample application, we identified several GWAS and eQTL variants with potential regulatory impact that also occur at the same base position as the rarer ClinVar variants (Figure 6D), which we suggest are ideal candidates for functional analysis. We highlight a few examples (Figure 6D) to illustrate this potential. In one case, a G>A substitution at chr21:44,886,223 in the *CD18/ITGB2* gene is a GWAS variant associated with neutrophil count. ITGB2 is an integrin important for cell surface adhesion, and biallelic mutations in this gene result in leukocyte adhesion deficiency, and increased neutrophil count^50^. A G>T in the same position is a VUS for a patient with leukocyte adhesion deficiency in ClinVar. These variants are in a FUS motif as well as a UPF1 eCLIP peak in HepG2 cells (Fig. 3A), suggesting potential regulatory impact. Another variant, a C>A at chr11:124,921,026 in the *HEPACAM* gene is an eQTL in nervous tissue, and is also located in EIF4G2 and FUS motifs, as well as a 7mer-m8 site for miR-6840-5p. A C>T in the same position is a VUS in ClinVar for megalencephalic leukoencephalopathy, which results from biallelic mutations in *HEPACAM*^51^. These variants should be investigated as potentially pathogenic for these patients, as GWAS or eQTL data support their causality, and RegVar supports the regulatory activity of these variants, suggesting possible mechanisms of action.

## DISCUSSION

Characterization of noncoding variants is required to expand the impact of exome and genome sequencing on the clinical sphere. Here, we show that 3’UTR variants in eCLIP peaks, RBP motifs, miRNA seed sites, and common APA isoform regions proximal to polyA sites are associated (often strongly) with gene expression changes, phenotypes, and pathology. We provide a tool, RegVar, to help researchers and clinicians prioritize noncoding variants for functional analysis based on their location in regulatory elements. RegVar can process many variants in parallel and can be readily integrated into bioinformatic pipelines for systematic variant annotation. We anticipate that this program will be used to interpret the over 10,000 ClinVar variants of uncertain significance in putative 3’UTR regulatory elements.

New models are being developed to predict whether variants have *cis*-regulatory effects on gene expression or phenotype^52–55^. These models incorporate many variant annotations, but have not incorporated RBP motif, miRNA target, or eCLIP peak data. Our general linearized model suggests that up to 10% more high-confidence GWAS hits or eQTL variants can be explained with incorporation of these features. Our findings will improve discrimination of pathogenic variants, as we show that 3’UTR variants with the same CADD score are more likely to affect gene expression or phenotype if they fall in these regulatory elements. In addition, our tool, RegVar, fills an important unmet need in noncoding variant interpretation, providing variant effect prediction.

Recent findings show that eQTLs are mostly found in less constrained genes with simple regulatory architecture, compared to GWAS hits, which are more likely to be found in functionally-important genes^47^. This could suggest that predicting variants that impact gene expression has limited clinical utility. However, we found that eQTL variants in regulatory elements are more likely to be GWAS hits, indicating that including regulatory features into eQTL models will help distinguish phenotypically-important eQTLs.

Most patients with undiagnosed rare diseases have exome rather than genome sequencing performed despite the increased power of genome sequencing^56^. This is largely due to limitations on interpretation of noncoding variants^57^. Our findings argue that clinical sequencing should extend further into the 3’UTR to improve pathogenic variant detection. This could be done by extending exome capture slightly without significantly increasing the cost of sequencing. Recently published guidelines for interpretation of noncoding variants require querying multiple databases^58^. We propose noncoding variants should be systematically assessed using RegVar and reported with sequencing results. RegVar will decrease the workload for noncoding variant interpretation by incorporating multiple datasets into one user-friendly tool.

Despite the advances our study makes into interpretation of noncoding variants, there remain some limitations. Our findings are based on GWAS and eQTL variants, which are more common in the population than disease-causing variants. In addition, GWAS hits and eQTL variants are mostly of low PIP after fine-mapping, suggesting that most variants in these datasets are not truly causal. However, we found that pathogenic ClinVar variants, like eQTLs and GWAS hits, are more likely to be in RBP motifs and eCLIP peaks, suggesting our results are generalizable to rare, pathogenic variants. We anticipate that our work will result in targeted experimental studies of patient variants that will aid in disease diagnosis.

Our data provide a thorough analysis of 3’UTR variants from several computational and population-wide datasets. We found variants that exhibit allele-specific binding in cells are more likely to be in predicted motifs, suggesting these computational methods predict in vivo regulation. We limited our study to the 3’UTR because this is an important regulatory region ignored by current methods of variant effect prediction; however, RBPs, and to a more limited extent miRNAs, also bind other noncoding regions such as introns or 5’UTR. Our results may extend to these areas as well, and these will be important future areas of study as variants in these regions also remain difficult to interpret.

## ACKNOWLEDGEMENTS

We thank members of the Burge laboratory for their helpful discussions and comments on the manuscript, especially Michael McGurk and Hannah Jacobs for their insights. In addition, we thank Sarah Stenton and Anne O’Donnell Luria for their helpful comments on the manuscript. L. Romo was supported by a Fred Lovejoy Resident Research and Education Fund award. This work was funded by grants from the NIH (GM085319 and HG002439 to C.B.B.).

## AUTHOR CONTRIBUTIONS

L.R. designed the study with input from C.B.B. and S.D.F. L.R. performed analysis with contributions from S.D.F.; L.R. wrote the RegVar package; L.R. wrote the draft manuscript; L.R. and C.B.B. finalized the manuscript with input from S.D.F.

## DATA AND CODE AVAILABILITY

The fine-mapped eQTL and GWAS datasets analyzed during the current study are publicly available from the GTEx repository and Finucane lab website: https://gtexportal.org/home/datasets, https://www.finucanelab.org/data. Allele specific eCLIP binding events were downloaded from from the Xiao lab supplemental data https://www.nature.com/articles/s41467-019-09292-w#Sec18, and all possible heterozygous background variants were kindly provided by the Xiao lab upon request.

## SUPPLEMENTAL FIGURES

**Figure S1:**
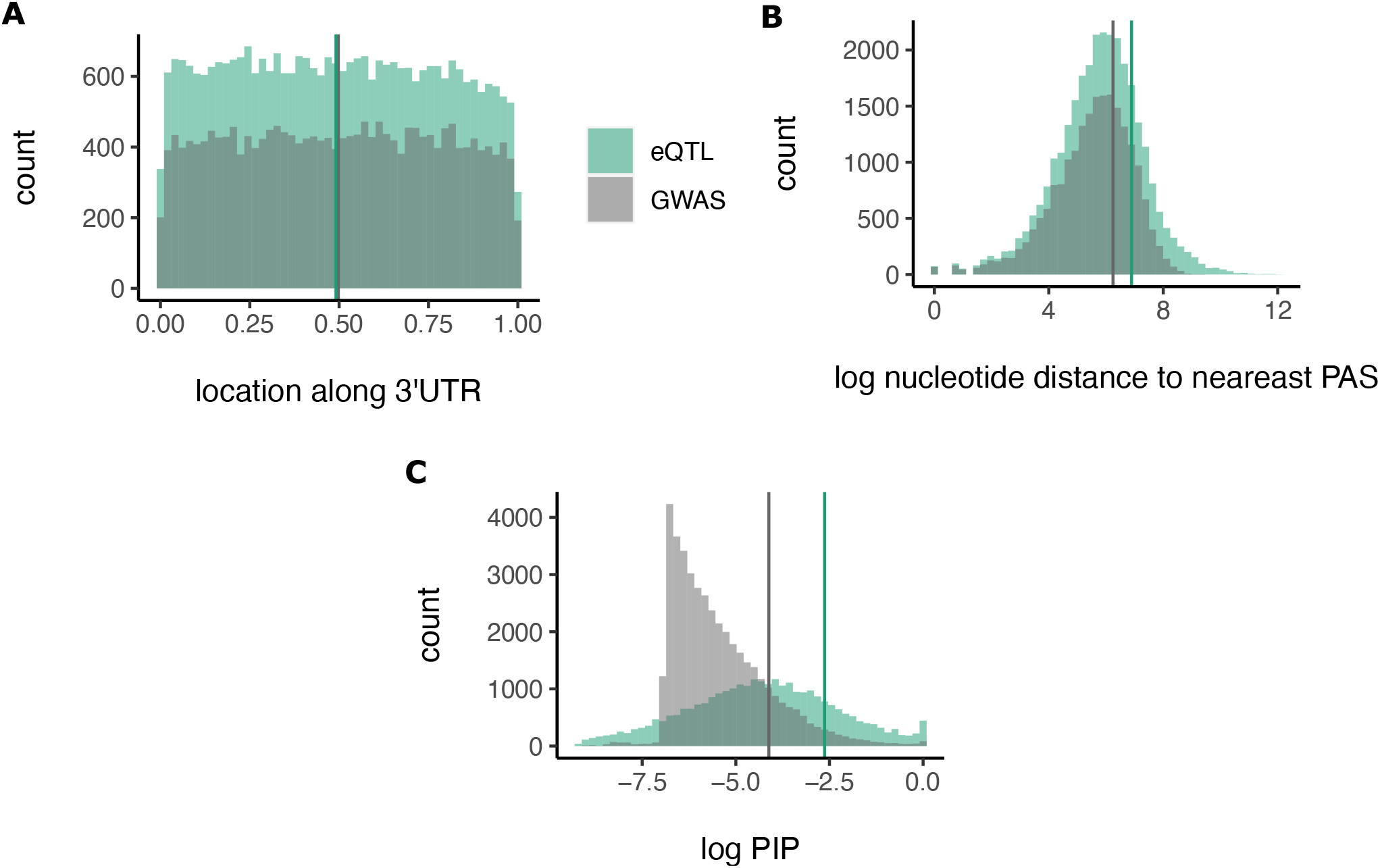
Comparison of eQTL variant and GWAS hit distribution along 3’UTR (A), proximity to nearest polyA signal (B), and PIPs (C).

**Figure S2:**
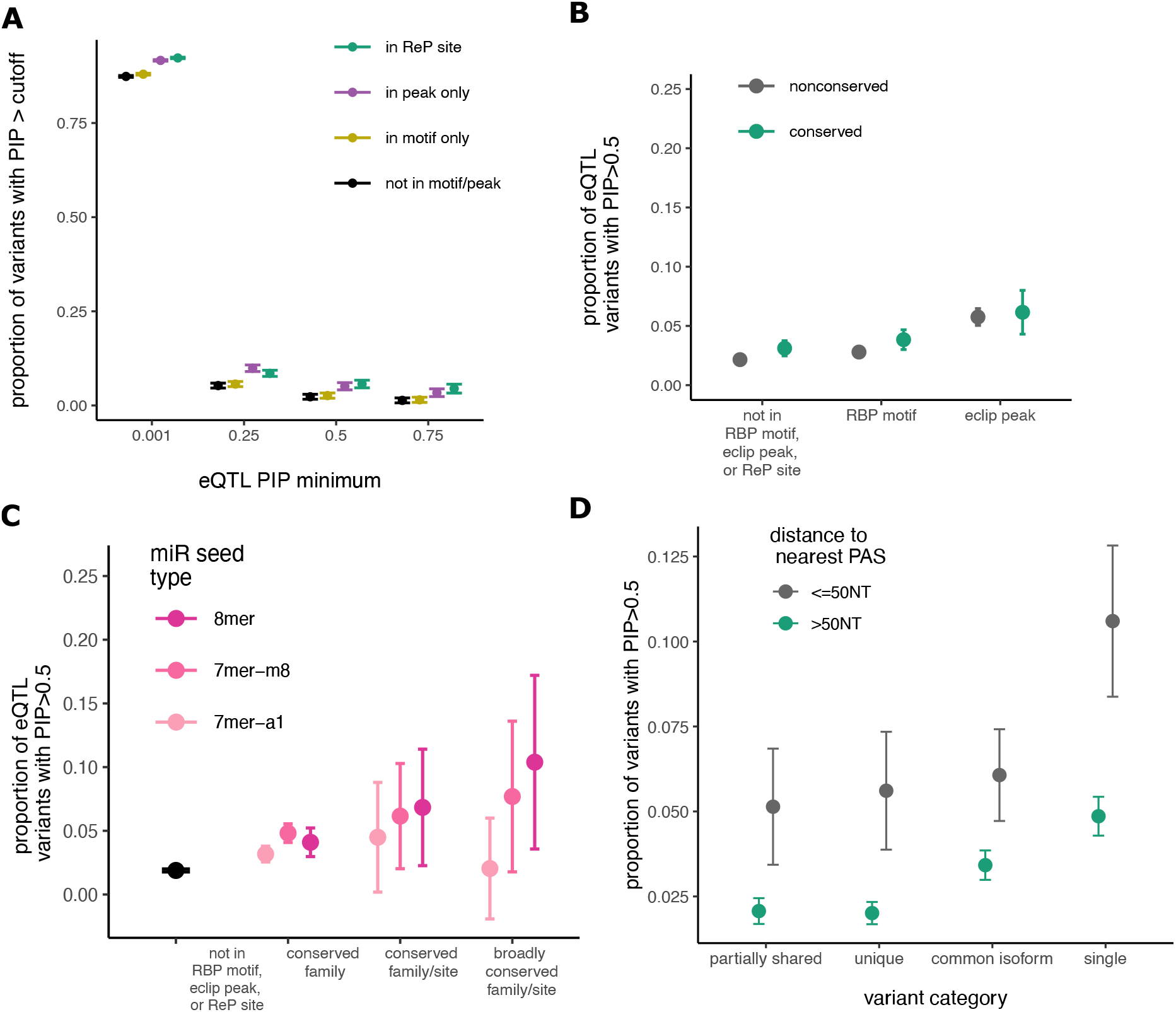
eQTL findings are robust even with a more stringent summary statistic PIP threshold. **A.** Proportion of eQTLs with PIP greater than a minimum cutoff for variants not in RBP motifs or eCLIP peaks compared to variants in RBP motifs, eCLIP peaks, and ReP sites, with 95% confidence intervals. **B.** Fraction causal (proportion of eQTLs with PIP>0.5) for variants not in RBP motifs or eCLIP peaks compared to variants in RBP motifs or eCLIP peaks. **C.** Fraction causal for variants not in miRNA sites compared to variants in miRNA sites with increasing predicted seed strength. **D.** Fraction causal for eQTL variants in genes with various numbers of canonical alternatively polyadenylated (APA) isoforms.

**Figure S3:**
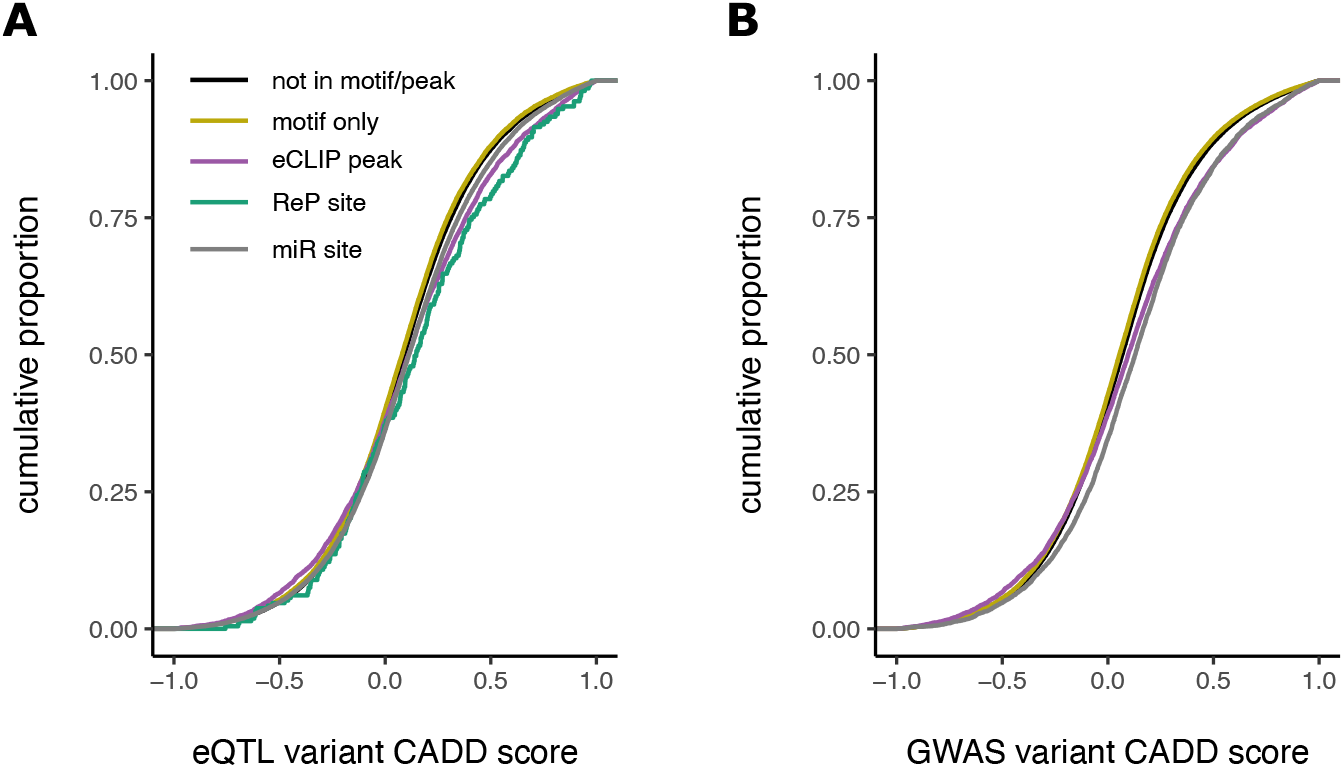
Variants in putative regulatory elements have higher CADD scores. Comparison of raw combined annotation dependent depletion (CADD) score distributions for eQTLs (A) or GWAS hits (B) in various putative regulatory elements versus controls.

**Figure S4:**
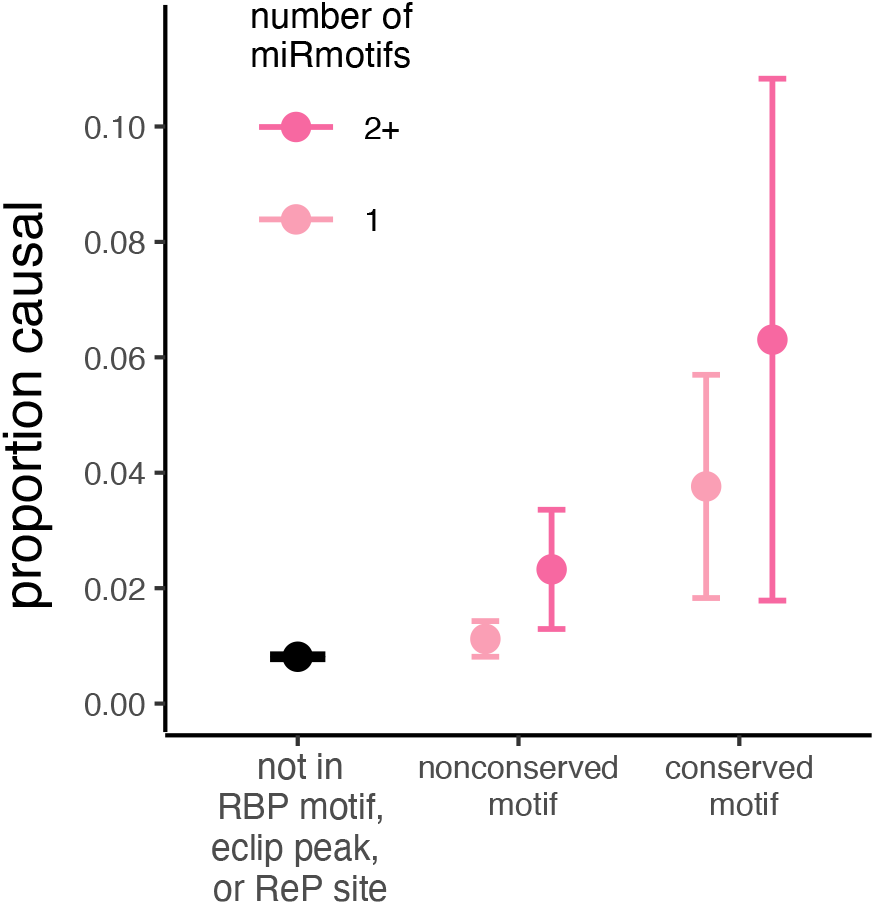
Trend towards higher PIP for variants predicted to disrupt more than one miRNA site. Fraction causal for GWAS variants not in miRNA sites compared to variants in increasing number of sites.

**Figure S5:**
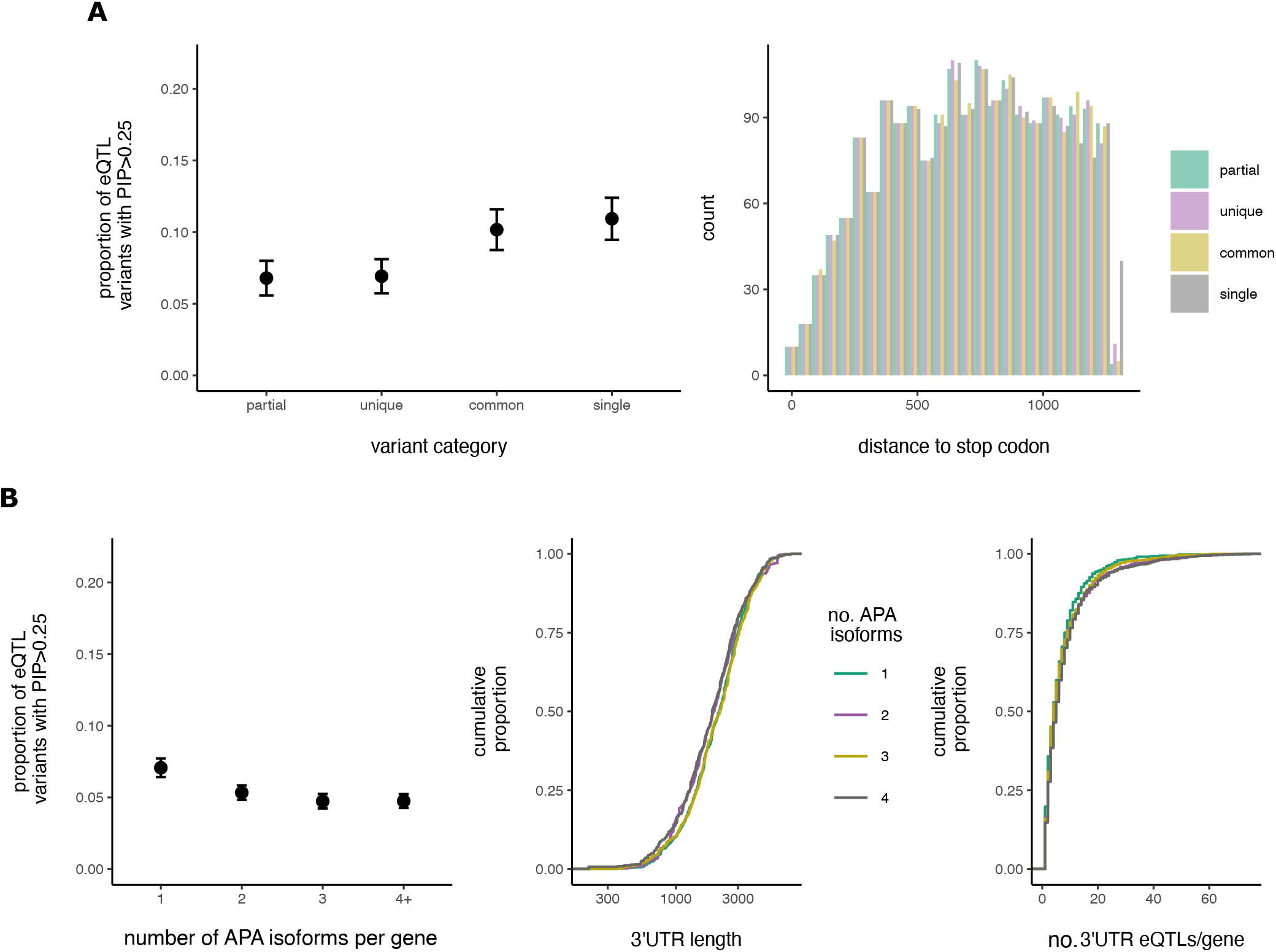
eQTL findings are not due to stop proximity, 3’UTR length, or number of eQTLs per gene. **A.** Fraction causal (proportion of eQTL variants with PIP greater than 0.25) for variants in various 3’UTR regions (left), after matching distance to canonical stop codon (right). **B.** Fraction causal for eQTL variants in genes with various numbers of canonical alternatively polyadenylated (APA) isoforms (left) after matching gene 3’UTR length (middle). On right is the distribution of number of eQTLs per gene for genes with varying isoform numbers.

**Figure S6:**
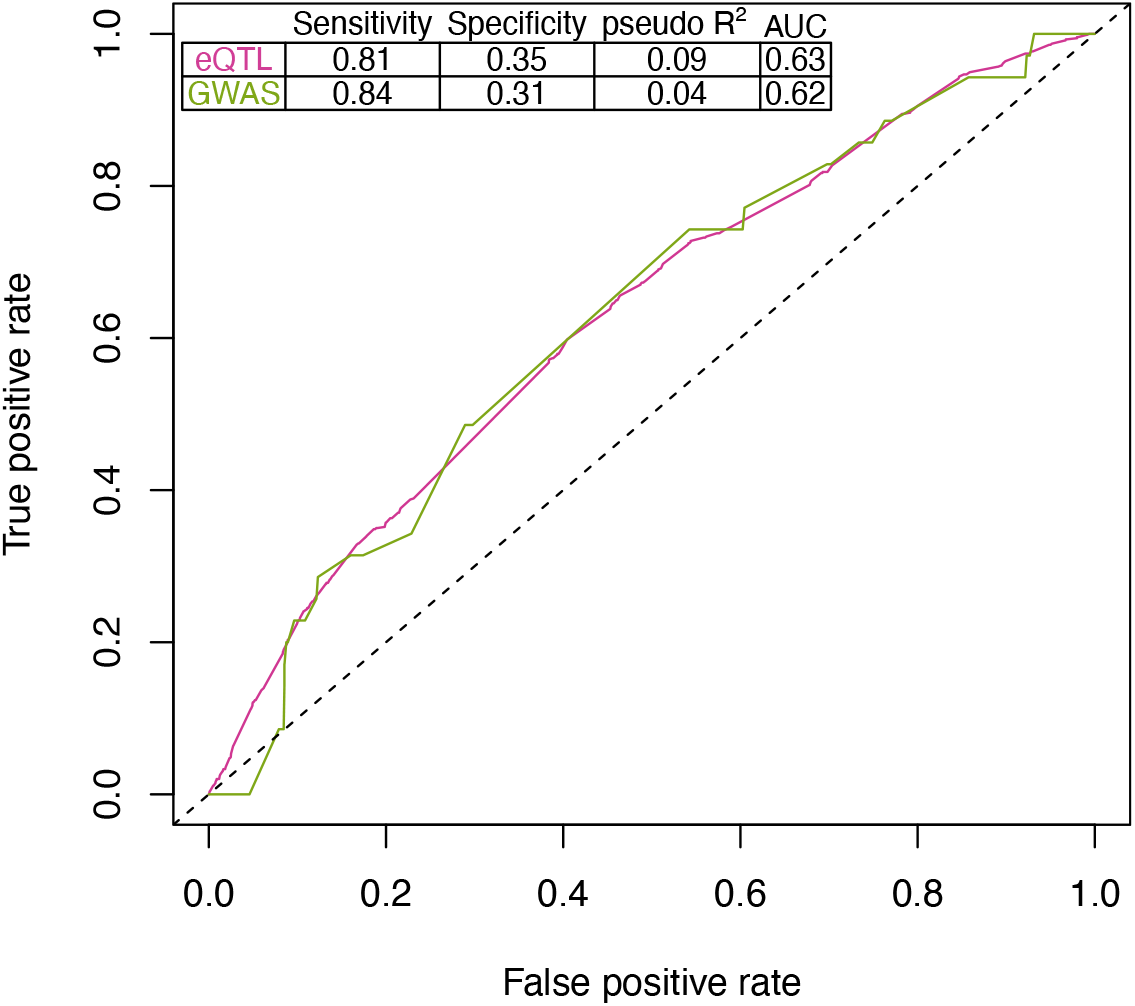
Performance of generalized linear models. Logistic regression analysis was performed to predict GWAS and eQTL variants (PIP>0.5). A variant was predicted to be an eQTL or GWAS hit if its log-odds was greater than 0.01 (eQTL) or 0.0075 (GWAS). These thresholds maximized sensitivity and specificity. Goodness of fit was assessed via Hosmer-Lemeshow Test with a chi squared of 1.0204 and p-value of 0.9981 for the eQTL model and a chi squared of 13.262 and p-value of 0.1032 for the GWAS model.

**Figure S7:**
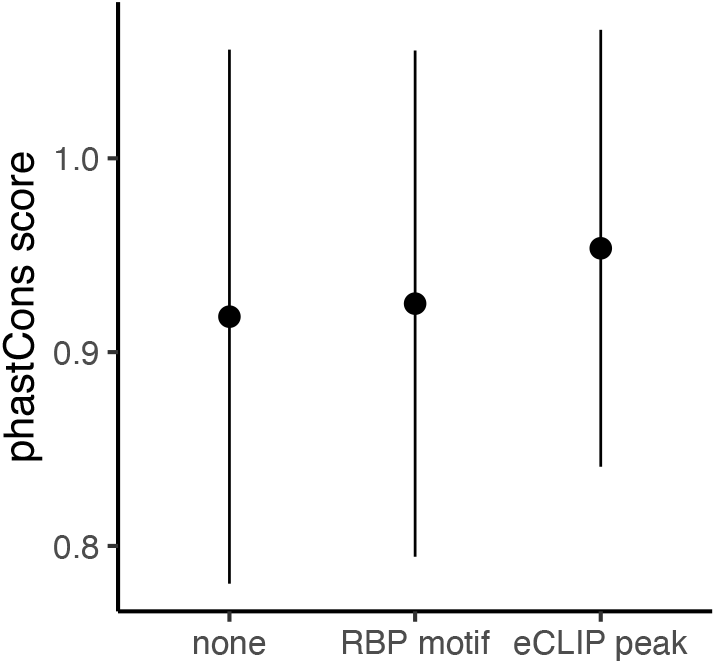
enrichment for pathogenic variants in regulatory elements is not solely due to conservation. Shown is mean phastCons score with standard deviation for variants in each category.

## REFERENCES

1. Dewey, F. E. et al. Distribution and clinical impact of functional variants in 50,726 whole-exome sequences from the DiscovEHR study. Science (80-.). 354, (2016).

2. Ng, P. C. & Henikoff, S. SIFT: Predicting amino acid changes that affect protein function. Nucleic Acids Res. 31, 3812–3814 (2003).

3. Adzhubei, I. A. et al. A method and server for predicting damaging missense mutations. Nature Methods (2010) doi:10.1038/nmeth0410-248.

4. Karczewski, K. J. et al. Nature-2020-Karczewski-MacArthur-The mutational constraint spectrum quantified from variation in 141,456 humans_supplement.pdf.

5. Pérez-Palma, E., Gramm, M., Nürnberg, P., May, P. & Lal, D. Simple ClinVar: an interactive web server to explore and retrieve gene and disease variants aggregated in ClinVar database. Nucleic Acids Res. 47, W99–W105 (2019).

6. Gebauer, F., Schwarzl, T., Valcárcel, J. & Hentze, M. W. RNA-binding proteins in human genetic disease. Nat. Rev. Genet. 22, 185–198 (2021).

7. Mayya, V. K. & Duchaine, T. F. Ciphers and executioners: How 3 0 -untranslated regions determine the fate of messenger RNAs. Front. Genet. 10, 1–18 (2019).

8. Tian, B. & Manley, J. L. Alternative polyadenylation of mRNA precursors. Nat. Rev. Mol. Cell Biol. 18, 18– 30 (2016).

9. Jiang, P. & Coller, H. Functional Interactions Between microRNAs and RNA Binding Proteins. MicroRNA e (2012) doi:10.2174/2211536611201010070.

10. Gebert, L. F. R. & MacRae, I. J. Regulation of microRNA function in animals. Nature Reviews Molecular Cell Biology (2019) doi:10.1038/s41580-018-0045-7.

11. Lambert, N. et al. RNA Bind-n-Seq: Quantitative Assessment of the Sequence and Structural Binding Specificity of RNA Binding Proteins. Mol. Cell 54, 887–900 (2014).

12. Van Nostrand, E. L. et al. Robust transcriptome-wide discovery of RNA-binding protein binding sites with enhanced CLIP (eCLIP). Nat. Methods 13, 508–514 (2016).

13. Agarwal, V., Bell, G. W., Nam, J. W. & Bartel, D. P. Predicting effective microRNA target sites in mammalian mRNAs. Elife 4, 1–38 (2015).

14. Fields, C. J. et al. Sequencing of Argonaute-bound microRNA/ mRNA hybrids reveals regulation of the unfolded protein response by microRNA-320a. PLoS Genet. 17, 1–28 (2021).

15. Rentzsch, P., Witten, D., Cooper, G. M., Shendure, J. & Kircher, M. CADD: Predicting the deleteriousness of variants throughout the human genome. Nucleic Acids Res. 47, D886–D894 (2019).

16. Abascal, F. et al. Expanded encyclopaedias of DNA elements in the human and mouse genomes. Nature 583, 699–710 (2020).

17. Ionita-Laza, I., Mccallum, K. & Buxbaum, J. A SPECTRAL APPROACH INTEGRATING FUNCTIONAL GENOMIC ANNOTATIONS FOR CODING AND NONCODING VARIANTS IULIANA IONITA-LAZA HHS Public Access Author manuscript. Nat Genet 48, 214–220 (2016).

18. Quang, D., Chen, Y. & Xie, X. DANN: A deep learning approach for annotating the pathogenicity of genetic variants. Bioinformatics 31, 761–763 (2015).

19. Hassan, M. S., Shaalan, A. A., Dessouky, M. I., Abdelnaiem, A. E. & ElHefnawi, M. A review study: Computational techniques for expecting the impact of non-synonymous single nucleotide variants in human diseases. Gene 680, 20–33 (2019).

20. Frazer, J. et al. Disease variant prediction with deep generative models of evolutionary data. Nature 599, 91–95 (2021).

21. Mohammadi, P., Castel, S. E., Brown, A. A. & Lappalainen, T. Quantifying the regulatory effect size of cis-acting genetic variation using allelic fold change. Genome Res. 27, 1872–1884 (2017).

22. Bycroft, C. et al. The UK Biobank resource with deep phenotyping and genomic data. Nature 562, 203– 209 (2018).

23. Wen, X. et al. The UK10K project identifies rare variants in health and disease. Nature 10, 1–29 (2019).

24. Aguet, F. et al. Genetic effects on gene expression across human tissues. Nature 550, 204–213 (2017).

25. Griesemer, D. et al. Genome-Wide Functional Screen of 3’UTR Variants Uncovers Causal Variants for Human Disease and Evolution. SSRN Electron. J. 1–18 (2021) doi:10.2139/ssrn.3762769.

26. Klein, J. C. et al. Functional testing of thousands of osteoarthritis-associated variants for regulatory activity. Nat. Commun. 10, 1–9 (2019).

27. Findlay, G. M. et al. Accurate classification of BRCA1 variants with saturation genome editing. Nature 562, 217–222 (2018).

28. Meng, L. et al. Use of exome sequencing for infants in intensive care units ascertainment of severe single-gene disorders and effect on medical management. JAMA Pediatr. 171, 1–10 (2017).

29. Krantz, I. D. et al. Effect of Whole-Genome Sequencing on the Clinical Management of Acutely Ill Infants with Suspected Genetic Disease: A Randomized Clinical Trial. JAMA Pediatr. 92122, 1218–1226 (2021).

30. Wen, X., Pique-Regi, R. & Luca, F. Integrating molecular QTL data into genome-wide genetic association analysis: Probabilistic assessment of enrichment and colocalization. PLoS Genet. 13, 1–25 (2017).

31. Wang, R., Nambiar, R., Zheng, D. & Tian, B. PolyA-DB 3 catalogs cleavage and polyadenylation sites identified by deep sequencing in multiple genomes. Nucleic Acids Res. (2018) doi:10.1093/nar/gkx1000.

32. Quinlan, A. R. & Hall, I. M. BEDTools: A flexible suite of utilities for comparing genomic features. Bioinformatics 26, 841–842 (2010).

33. Schaid, D. J., Chen, W. & Larson, N. B. From genome-wide associations to candidate causal variants by statistical fine-mapping. Nat. Rev. Genet. 19, 491–504 (2018).

34. Wen, X., Luca, F. & Pique-Regi, R. Cross-Population Joint Analysis of eQTLs: Fine Mapping and Functional Annotation. PLoS Genet. 11, 1–29 (2015).

35. Wang, G., Sarkar, A., Carbonetto, P. & Stephens, M. A simple new approach to variable selection in regression, with application to genetic fine mapping. J. R. Stat. Soc. Ser. B Stat. Methodol. (2020) doi:10.1111/rssb.12388.

36. Lianoglou, S., Garg, V., Yang, J. L., Leslie, C. S. & Mayr, C. Ubiquitously transcribed genes use alternative polyadenylation to achieve tissue-specific expression. Genes Dev. 27, 2380–2396 (2013).

37. Romo, L., Ashar-Patel, A., Pfister, E. & Aronin, N. Alterations in mRNA 3ʹUTR isoform abundance accompany gene expression changes in human Huntington’s disease brains. Cell Rep. (2017).

38. Siepel, A. et al. Evolutionarily conserved elements in vertebrate, insect, worm, and yeast genomes. Genome Res. (2005) doi:10.1101/gr.3715005.

39. Yang, E. W. et al. Allele-specific binding of RNA-binding proteins reveals functional genetic variants in the RNA. Nat. Commun. 10, (2019).

40. Kircher, M. et al. A general framework for estimating the relative pathogenicity of human genetic variants. Nat. Genet. 46, 310–315 (2014).

41. Felsenstein, J. & Churchill, G. A. A Hidden Markov Model approach to variation among sites in rate of evolution. Mol. Biol. Evol. (1996) doi:10.1093/oxfordjournals.molbev.a025575.

42. Yang, R. et al. La-Related Protein 4 Binds Poly(A), Interacts with the Poly(A)-Binding Protein MLLE Domain via a Variant PAM2w Motif, and Can Promote mRNA Stability. Mol. Cell. Biol. (2011) doi:10.1128/mcb.01162-10.

43. Yang, H., Duckett, C. S. & Lindsten, T. iPABP, an inducible poly(A)-binding protein detected in activated human T cells. Mol. Cell. Biol. (1995) doi:10.1128/mcb.15.12.6770.

44. Dominguez, D., et al. Sequence, Structure, and Context Preferences of Human RNA Binding Proteins. Mol. Cell 70, 854–867.e9 (2018).

45. Natarajan, K., Sundaramoorthy, A. & Shanmugam, N. HnRNPK and lysine specific histone demethylase-1 regulates IP-10 mRNA stability in monocytes. European Journal of Pharmacology vol. 920 (Elsevier B.V., 2022).

46. Liu, X. H. et al. Regulation and related mechanism of GSN mRNA level by hnRNPK in lung adenocarcinoma cells. Biol. Chem. 400, 951–963 (2019).

47. Mostafavi, H., Spence, J. P., Naqvi, S. & Pritchard, J. K. Limited overlap of eQTLs and GWAS hits due to systematic differences in discovery. bioRxiv Prepr. 1–39 (2022).

48. Zhang, J. et al. RADAR: annotation and prioritization of variants in the post-transcriptional regulome of RNA-binding proteins. Genome Biol. (2020) doi:10.1186/s13059-020-01979-4.

49. Seitz, H. Issues in current microRNA target identification methods. RNA Biol. 14, 831–834 (2017).

50. van de Vijver, E. et al. Hematologically important mutations: Leukocyte adhesion deficiency (first update). *Blood Cells*, Mol. Dis. (2012) doi:10.1016/j.bcmd.2011.10.004.

51. López-Hernández, T. et al. Mutant GlialCAM causes megalencephalic leukoencephalopathy with subcortical cysts, benign familial macrocephaly, and macrocephaly with retardation and autism. Am. J. Hum. Genet. (2011) doi:10.1016/j.ajhg.2011.02.009.

52. Wang, Q. S. et al. Leveraging supervised learning for functionally informed fine-mapping of cis-eQTLs identifies an additional 20,913 putative causal eQTLs. Nat. Commun. 12, 1–11 (2021).

53. Agarwal, V. & Shendure, J. Predicting mRNA Abundance Directly from Genomic Sequence Using Deep Convolutional Neural Networks. Cell Rep. 31, 107663 (2020).

54. Zhou, J. & Troyanskaya, O. G. Predicting effects of noncoding variants with deep learning-based sequence model. Nat. Methods 12, 931–934 (2015).

55. Zhou, J. et al. Deep learning sequence-based ab initio prediction of variant effects on expression and disease risk. Nat. Genet. 50, 1171–1179 (2018).

56. Belkadi, A. et al. Whole-genome sequencing is more powerful than whole-exome sequencing for detecting exome variants. Proc. Natl. Acad. Sci. U. S. A. 112, 5473–5478 (2015).

57. Petersen, B. S., Fredrich, B., Hoeppner, M. P., Ellinghaus, D. & Franke, A. Opportunities and challenges of whole-genome and -exome sequencing. BMC Genet. 18, 1–13 (2017).

58. Ellingford, J. M. et al. Recommendations for clinical interpretation of variants found in non-coding regions of the genome. Genome Med. (2022) doi:10.1186/s13073-022-01073-3.

